# Vinculin is Essential For Sustaining Normal Levels of Endogenous Force Transmission at Cell-Cell Contacts

**DOI:** 10.1101/2023.09.05.556369

**Authors:** Mazen Mezher, Sandeep Dumbali, Ian Fenn, Carter Lamb, Conrad Miller, Jolene I. Cabe, Vidal Bejar-Padilla, Daniel Conway, Venkat Maruthamuthu

## Abstract

Transmission of cell-generated (i.e., endogenous) tension at cell-cell contacts is crucial for tissue shape changes during morphogenesis and adult tissue repair in tissues like epithelia. E-cadherin-based adhesions at cell-cell contacts are the primary means by which endogenous tension is transmitted between cells. The E-cadherin-β-catenin-α-catenin complex mechanically couples to the actin cytoskeleton (and thereby the contractile machinery of the cell) both directly and indirectly. However, the key adhesion constituents required for substantial endogenous force transmission at these adhesions in cell-cell contacts are unclear. Due to the role of α-catenin as a mechanotransducer that recruits vinculin at cell-cell contacts, we expected α-catenin to be essential for the high levels of force transmission normally sustained. Instead, using the traction force imbalance method to determine the inter-cellular force at a single cell-cell contact between cell pairs, we found that it is vinculin that is essential for high endogenous force transmission. Our results constrain the potential mechanical pathways of force transmission at cell-cell contacts and suggest that vinculin can transmit forces at E-cadherin adhesions independent of α-catenin, possibly through β-catenin. Furthermore, we tested the ability of cell-cell contacts to withstand external stretch and found that vinculin is essential to maintain cell-cell contact stability under external forces as well.

## Introduction

Cell-cell contacts in tissues like epithelia are interfaces where cell-generated and external forces are transmitted from cell to cell and thereby across the tissue [1, 2]. During morphogenesis, cell-generated tension transmitted through cell-cell contacts is essential for cell shape changes as well as supracellular morphological transformations [3]. Even pathological events such as cancer metastasis and cell-to-cell transmission of some pathogens involve changes in the forces transmitted at cell-cell contacts. These cell-cell contacts are bound by many types of adhesions, but E-cadherin adhesions are chiefly important in the integrity and mechanical function of these cell-cell contacts [4]. While biophysical and biomimetic approaches have broadened our understanding of E-cadherin adhesions and their response to forces [5, 6], there is much unknown about endogenous force transmission through E-cadherin adhesions in the native context of lateral contacts between cells. It is also unclear as to what specific factors determine the adhesion strength of these lateral cell-cell contacts when subject to external forces.

E-cadherin forms a 1:1 complex with β-catenin which in turn binds to α-catenin [7]. This E-cadherin-β-catenin-α-catenin complex can couple to the actin cytoskeleton by directly binding to actin, with enhanced binding of α-catenin to F-actin under force [8-10]. The E-cad-catenin complex can also couple (or potentially couple) to actin via the adhesion-associated protein vinculin [11], actin cross-linker α-actinin [12], tight junction protein ZO-1 [13], F-actin binding proteins afadin [14] or EPLIN [15] and the formin Fmn1 [16]. In particular, α-catenin has been shown to function as an elastic link in series with cadherin and actin [17] that transitions to an open conformation under force that then recruits vinculin [18]. Accordingly, such recruitment of vinculin has been shown to depend on non-muscle myosin II activity in cells [19]. Vinculin can also be recruited to E-cadherin adhesions via myosin VI [20] as well as β-catenin dependent ways [11, 21, 22], suggesting that vinculin may play a significant role in cell-to-cell force transmission. However, how much vinculin affects the endogenous force transmission at cell-cell contacts between epithelial cells, and how this compares to the contribution of a presumably more constitutive component like α-catenin, has not been directly assessed.

The mechanical function of cell-cell adhesion associated proteins are at least two-fold: Transmission of mechanical forces from cell to cell as well as maintenance of the strength of these adhesions. Many of the approaches used to study the role of cell-cell adhesion proteins in force transmission employ biochemical methods at cell-cell contacts themselves, or quantitative methods with biomimetic interfaces such as cadherin-coated substrates [5, 23-30], cadherin-coated beads [31, 32] and suspended cell doublets [33]. Quantitative approaches such as FRET-based sensors can look at force transmission through specific proteins at cell-cell contacts [6, 34, 35], but the total forces transmitted via cell-cell contacts are not known in this context. Due to the presence of multiple adhesion systems at cell-cell contacts, determining the total endogenous force transmitted at cell-cell contacts as such can help identify the overall effect of perturbations like specific protein knockdowns. In a similar manner, while cadherin-coated substrates and suspended cell doublets have enabled key insights into determinants of E-cadherin adhesion strength, assessment of adhesion strength of lateral cell-cell contacts between epithelial cells is essential to understand how this interface ultimately resists mechanical challenges and the role played by specific cell adhesion associated proteins such as vinculin. Here, we test the importance of putative physical pathways of force transmission by knocking out proteins prominently known to be involved in mechanotransduction at E-cadherin adhesions. We find that, contrary to expectation, vinculin rather than α-catenin is crucial for transmitting high endogenous tension at cell-cell contacts. We also use large external stretching to find that vinculin is essential for maintaining cell-cell contact integrity under external stretch, highlighting the crucial mechanical role of vinculin at epithelial cell-cell contacts.

## Materials and Methods

### Cell Culture

Madin-Darby Canine Kidney (MDCK) II cells were cultured in Dulbecco’s modified Eagle’s medium (Corning Inc., Corning NY) containing 10% fetal bovine serum (Corning Inc., Corning NY), L-glutamine, and 1 % Penicillin/Streptomycin at 37°C, under 5 % CO_2._ MDCK cells were plated overnight onto collagen I-coated soft silicone atop 22 mm square No.1.5 coverslips in 35 mm culture dishes and then used for experiments.

To generate knockout (KO) cells, CRISPR/Cas9 was used with the gRNA sequence CACGAGGAAGGCGAGGTGGA for vinculin (previously shown [36] to knockout vinculin) and the gRNA sequence TCTGGCAGTTGAAAGACTGT for α-catenin (previously shown [36] to knockout α-catenin). The gRNA sequences were used in the Sigma All-in-One U6-gRNA/CMV-Cas9-tGFP Vector). Cells were transiently transfected with this vector (with the appropriate gRNA), followed by clonal expansion. Clones were screened for vinculin or α-catenin loss using Western blotting. For the double (α-catenin and vinculin) KO, vinculin KO cells were used to generate an additional KO of α-catenin. α-catenin with its vinculin binding site (amino acids 316-405 in α-catenin) replaced by a homologous similar sequence from vinculin (amino acids 514-606 in vinculin), called α-catenin DVBS (delta vinculin binding site) [17, 37] was used to generate the α-catenin DVBS in α-catenin KO cell line (Addgene plasmid 178649).

### Western Blotting

Cells were washed with phosphate-buffered saline (PBS) and lysed using RIPA buffer supplemented with protease as well as phosphatase inhibitors. SDS–PAGE of the proteins was followed by Western blotting using PVDF membranes. The proteins on the PVDF membranes were incubated in 5% bovine serum albumin (BSA) in PBS for 1 h at room temperature followed by primary antibody incubation overnight. Then, the samples were rinsed in 0.2% Tween in PBS and incubated for 45 min with HRP-conjugated secondary antibody in 0.2% Tween in PBS. After rinsing with PBS, blot chemiluminescence was imaged using a Bio-Rad ChemiDoc system. Primary antibodies used for blotting were anti-rabbit α-catenin (catalogue# C2081 from Sigma, St. Louis, MO) vinculin (clone hVIN-1 from Sigma, St. Louis, MO), and anti-mouse tubulin (clone DM1A from Cell Signaling, Danvers, MA).

### Live Cell Imaging and Immunofluorescence

Leica DMi8 epifluorescence microscope (Leica Microsystems, Buffalo Grove, IL) was used to image live and fixed cells. An airstream incubator (Nevteck, Williamsville, VA) was used to maintain the temperature at 37°C during live cell imaging. Images were taken using a 40x objective lens and Clara cooled CCD camera (Andor Technology, Belfast, UK). MDCK cells were fixed utilizing 4% paraformaldehyde (Electron Microscopy Sciences, Hatfield, PA) in 1.5% Bovine Serum Albumin and 0.5% Triton. The actin cytoskeleton was stained using Alexa-488 Phalloidin from Thermo Fisher Scientific (Eugene,OR). Antibodies used were rabbit anti-vinculin (Abcam, catalog# ab129002), mouse anti-vinculin (clone hVIN-1, Sigma, Catalog #V9131), rabbit anti-α-catenin (Sigma, catalogue# C2081), mouse anti β-catenin (BD transduction laboratories, catalogue# 610153) and rabbit anti-E-cadherin (clone 24E10, Cell Signaling, Catalogue #3195S). Fluorophore-conjugated secondary antibodies from Jackson ImmunoResearch or ThermoFisher were used in all staining experiments.

### Preparation of Soft Silicone Substrate

Soft silicone (Qgel 300, CHT USA Inc., Richmond, VA) was prepared by mixing its A and B components at a 1:2.2 ratio. The gel mixture was cured using a heater at 100^0^ C for an hour. To use these silicone substrates for traction force microscopy, fluorescent beads and collagen I were coupled as follows. After curing, the silicone was exposed to 305 nm UV light (UVP cross-linker, Analytik Jena AG, Upland, CA) for 5 min. Red fluorescent beads of 0.44 µm diameter (with surface carboxyl groups) were coupled to the top surface of the silicone by incubating with an aqueous solution with 10 mg/mL EDC (1-ethyl-3-(3-dimethylaminopropyl) carbodiimide hydrochloride), 5 mg/mL sulfo-NHS (N-hydroxysulfosuccinimide) and 0.017 mg/mL collagen I for 30 minutes. Then, the substrate was washed with Dulbecco’s Phosphate-Buffered Saline (DPBS) before plating cells on it.

### Rheology of Soft Silicone

The shear rheology of the soft silicone was characterized using an MCR-302 rheometer (Anton Paar, Ashland, VA). Presence of an air bearing in the rheometer enabled measurement of the moduli of soft samples (kPa and below). The soft silicone was prepared and cured as above and loaded between 25 mm diameter parallel plates. The storage and loss shear moduli were obtained as a function of angular frequency for 1% strain (determined to be in the linear range using a strain sweep). The average of the shear storage modulus (G’) in the 0.1 to 1 rad/s range was considered to be the nominal G’.

### Traction Force Microscopy and Traction Force Imbalance Method

A phase image of each MDCK cell or cell pair along with the correspondent image of beads beneath were first recorded. After the cells were disintegrated using 1% sodium dodecyl sulfate, an image of the beads on the relaxed substrate were recorded. The stressed substrate bead images (in the presence of cells) and the relaxed bead images (in the absence of cells) were aligned using an ImageJ plugin [38]. The displacement field was then computed using mpiv (https://www.mathworks.com/matlabcentral/fileexchange/2411-mpiv), scripted in MATLAB (MathWorks, Natick, MA). Traction stresses are then reconstructed using regularized Fourier Transform Traction Cytometry that uses the Boussinesq solution, such as in previously published work [38-44]. The Traction Force Imbalance Method [26, 39, 41] was then used to compute the inter-cellular force at the cell-cell contact within a cell pair from the vector sum of traction forces under each cell within the cell pair.

### Biaxial Stretch of Epithelial Islands

A 0.01” thick silicone sheet (Speciality Manufacturing, Saginaw, MI) was exposed to 305 nm UV light for five minutes and then incubated with collagen I at 37^0^C, under 5% CO_2_ for 15 minutes. The sheet was washed with Dulbecco’s Phosphate-Buffered Saline (DPBS) and then MDCK cells were plated on the silicone sheet. After overnight culture, the cell culture medium was replaced with CellBrite (Biotium, Fremont, CA) in cell culture medium (used at 1:200) for 30 minutes at 37^0^C. Then, the silicone sheet (with cells) was placed inside the well of a custom-built biaxial cell stretcher. A phase and a fluorescence image (corresponding to plasma membrane staining with CellBrite) of the cells were taken before and after applying 2, 6, 15, 23 and 38% linear strain to the sheet.

### Statistical Analysis

For statistical analysis, t-test was used to compare wildtype and vinculin KO single cell data (fig. 1 and 5) and analysis of variance (ANOVA) was used for multiple comparisons of all the cell pair data (fig. 2, 3 and 4), with * indicating p < 0.05, ** indicating p < 0.01 and *** indicating p < 0.001.

**Figure 1.**
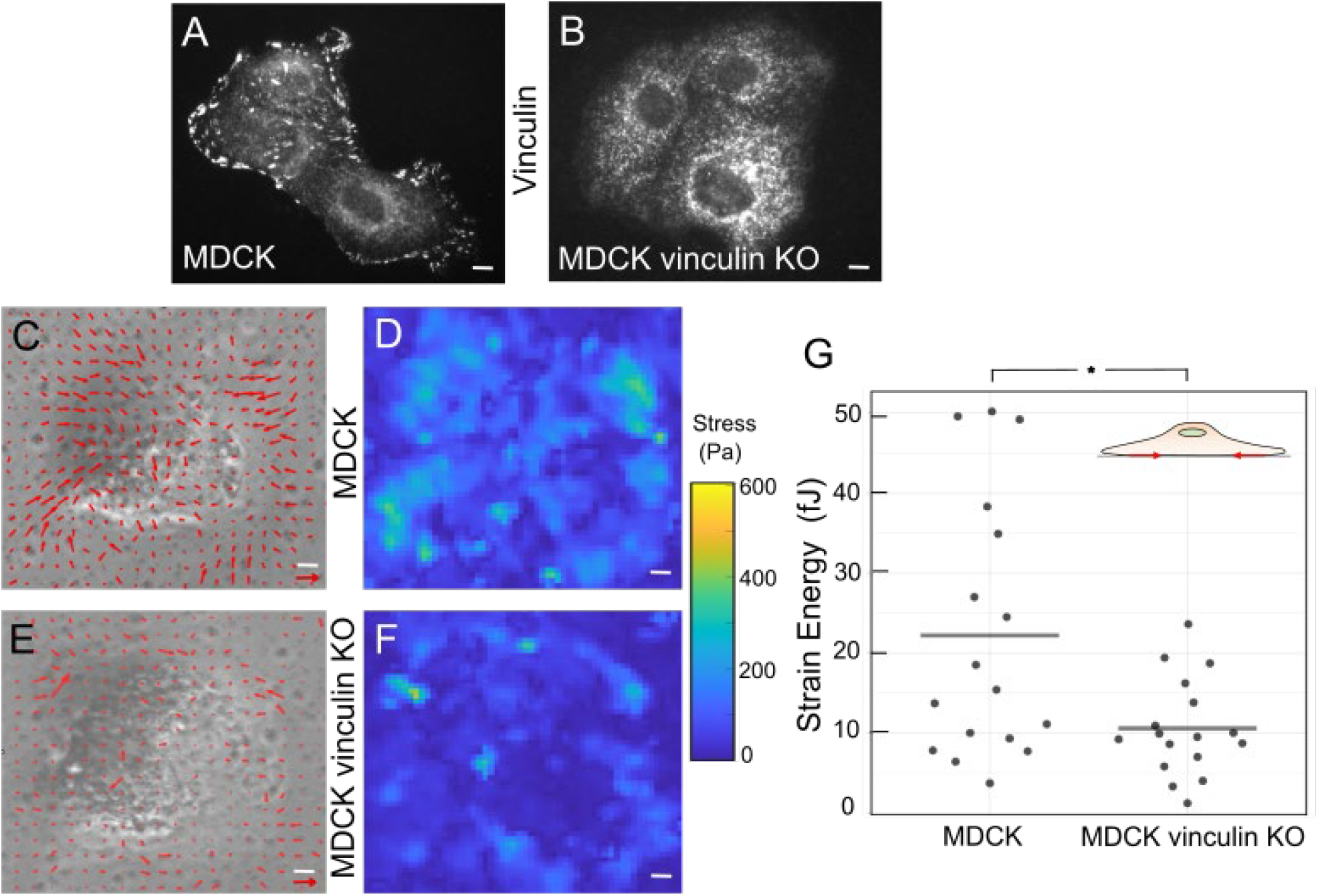
Loss of vinculin decreases cell-ECM traction forces. (A, B) Immunofluorescence images of MDCK (A) and MDCK vinculin KO (B) cells stained for vinculin. (C-F) Traction stresses exerted by a single MDCK cell (C) and MDCK vinculin KO (E) on the substrate. Traction stress vectors are overlaid (red arrows). Heat map of the traction magnitude for a single MDCK cell (D) and MDCK vinculin KO (F). Length scale bar (white) is 5 µm. Traction vector scale (red) is 400 Pa. (G) Plot of the strain energy due to traction force exertion for MDCK and MDCK vinculin KO single cells. Grey horizontal line represents the mean value. Inset shows schematic of a single cell to indicate that the plot data corresponds to traction forces exerted by single cells.

**Figure 2.**
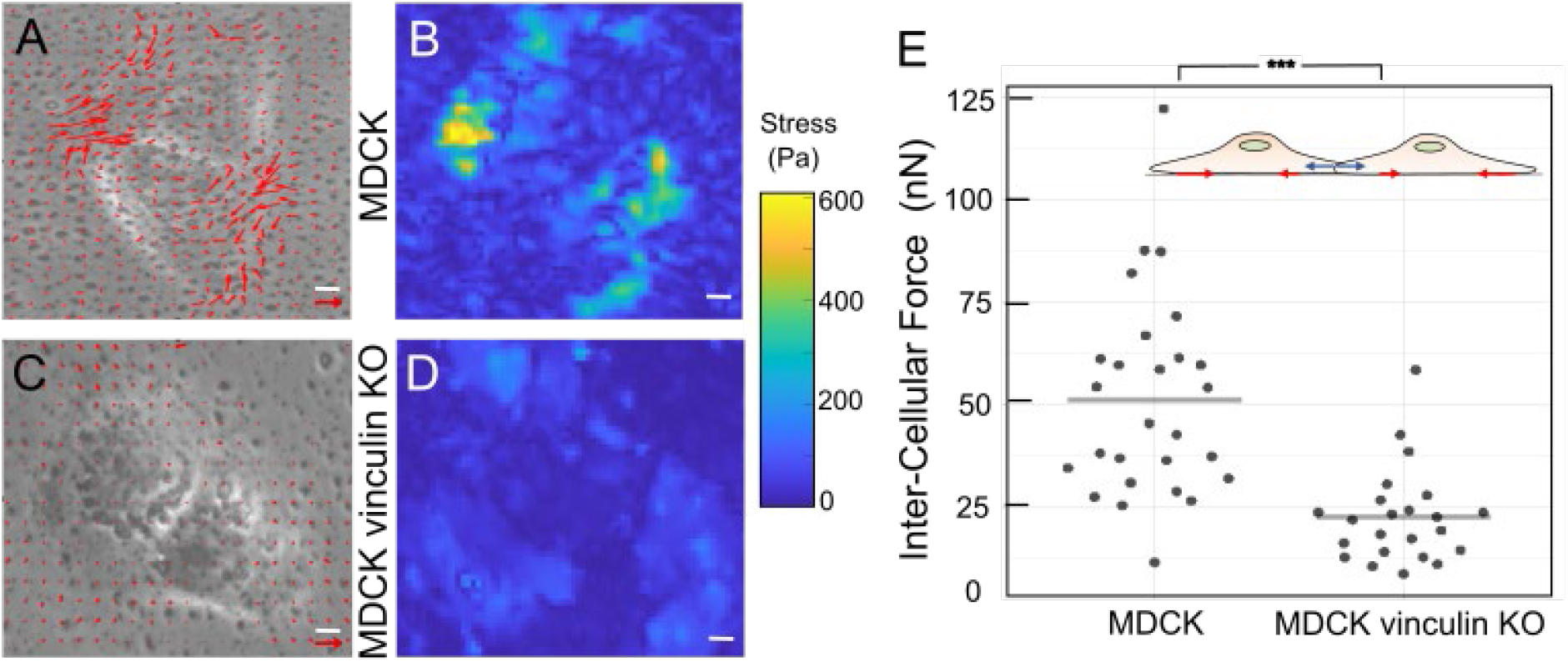
Loss of vinculin leads to a severe decrease in endogenous inter-cellular forces. (A-D) Traction stresses exerted by MDCK cell pair (A) and MDCK vinculin KO cell pair (C). Traction stress vectors are overlaid (red arrows.). Heat map of the traction magnitude for MDCK cell pair (B) and MDCK vinculin KO (D). Length scale bar (white) is 5 µm. Traction vector scale (red) is 400 Pa. (E) Plot of the intercellular forces for MDCK and MDCK vinculin KO cell pairs. Grey horizontal line represents the mean value. Inset shows schematic of a cell pair – the red traction vectors correspond to data in (A, C) and the blue inter-cellular force vectors have their magnitude plotted in (E).

**Figure 3.**
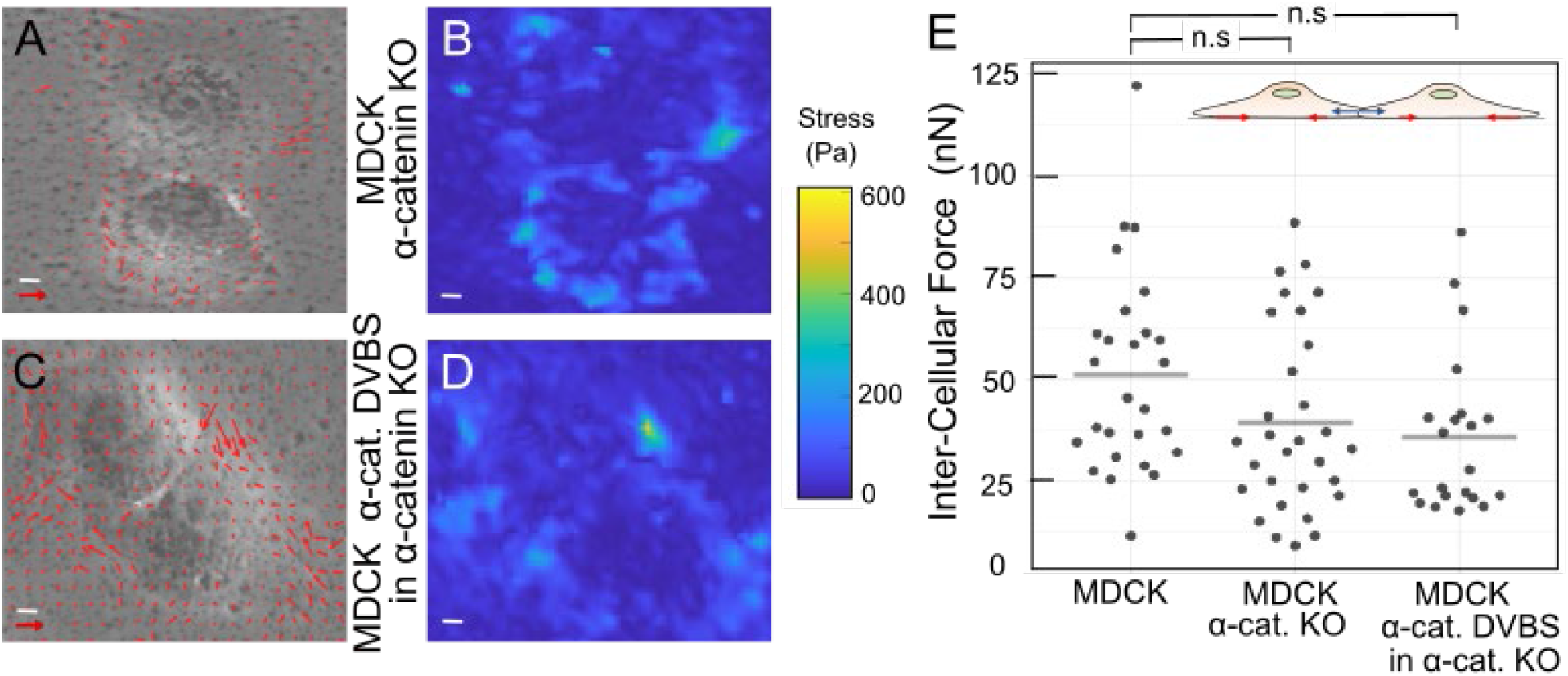
Loss of α-catenin does not significantly decrease inter-cellular forces. (A-D) Traction stresses of MDCK α-catenin cell pair (A) and MDCK α-catDVBS in α-catenin KO cell pair (C). Traction stress vectors are overlaid (red arrows.). Heat map of the traction magnitude for MDCK α-catenin KO cell pair (B) and MDCK α-catDVBS in α-catenin KO (D). Length scale bar (white) is 5 µm. Traction vector scale (red) is 400 Pa. (E) Plot of the intercellular forces for MDCK α-catenin KO and MDCK α-catenin DVBS in α-catenin KO cell pair. Black bar represents the mean value. Grey horizontal line represents the mean value. Inset shows schematic of a cell pair – the red traction vectors correspond to data in (A, C) and the blue inter-cellular force vectors have their magnitude plotted in (E). MDCK cell pair data in (E) is same as in Fig. 2E, reproduced here to enable comparison.

**Figure 4.**
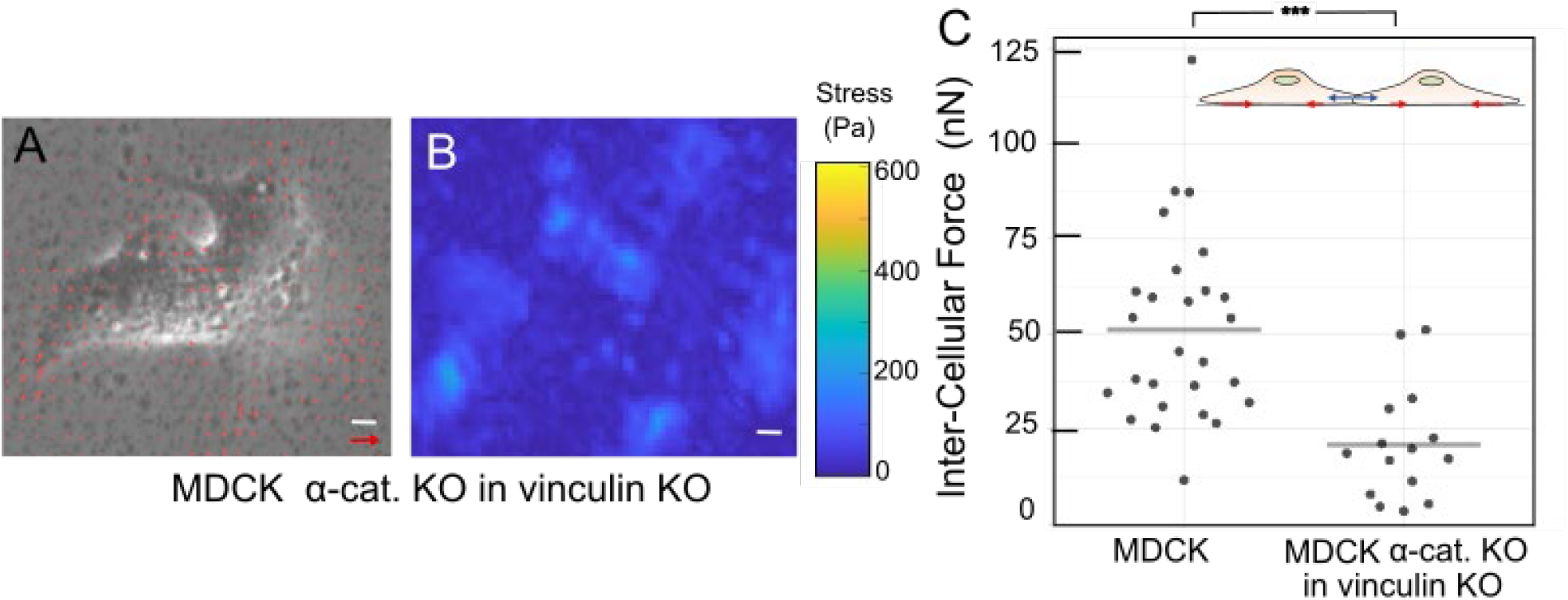
Loss of both α-catenin and vinculin leads to a severe decrease in inter-cellular forces. (A, B) Traction stresses of MDCK α-catenin KO in vinculin KO cell pair (A). Traction stress vectors are overlaid (red arrows.). Heat map of the traction magnitude for MDCK α-catenin KO in vinculin KO cell pair (B). Length scale bar (white) is 5 µm. Traction vector scale (red) is 400 Pa. (C) Plot of the intercellular forces for MDCK and MDCK α-catenin & vinculin KO cell pairs. Grey horizontal line represents the mean value. Inset shows schematic of a cell pair – the red traction vectors correspond to data in (A) and the blue inter-cellular force vectors have their magnitude plotted in (C). MDCK cell pair data in (C) is same as in Fig. 2E, reproduced here to enable comparison.

**Figure 5.**
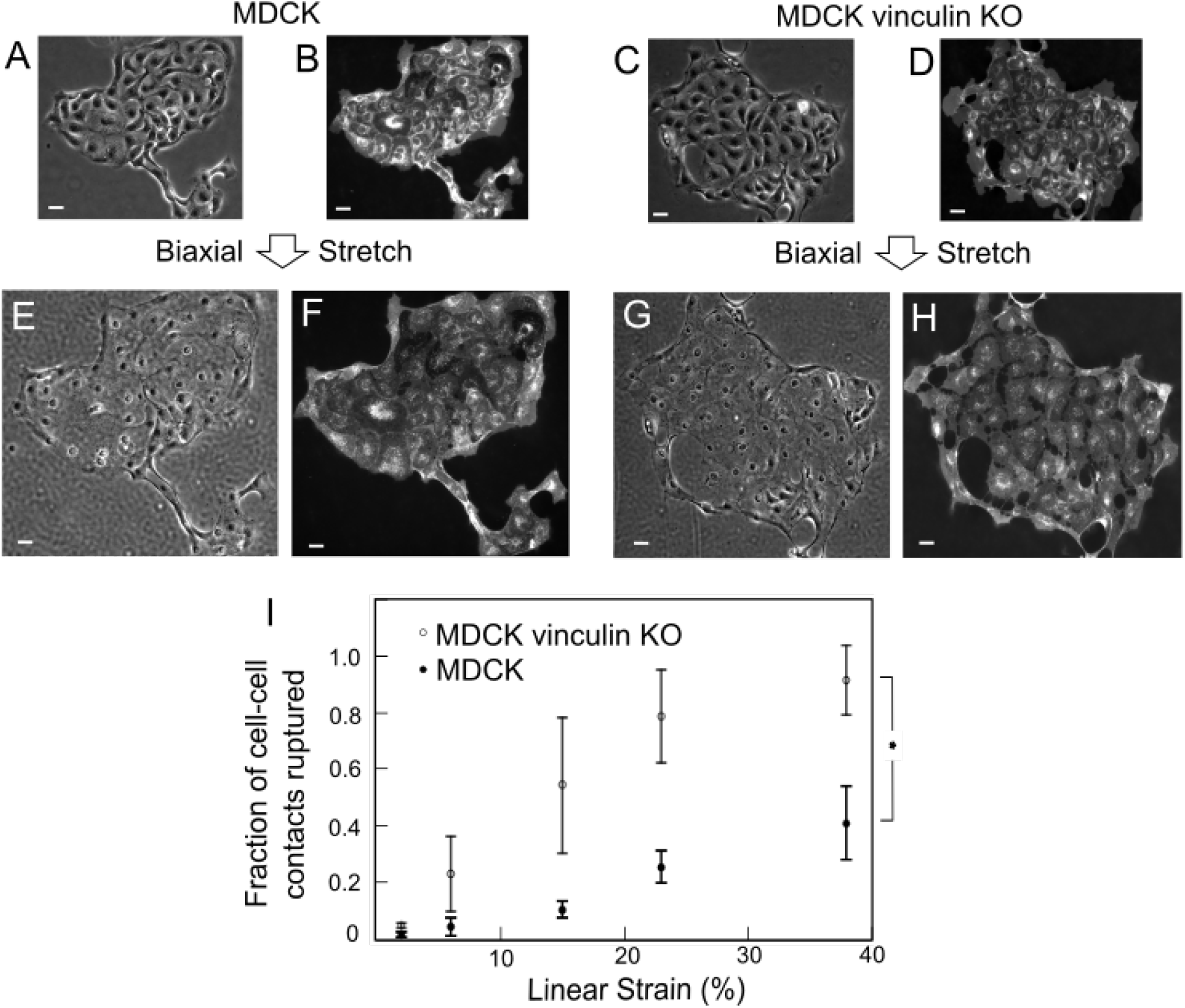
Vinculin is essential for the high adhesion strength of native cell-cell contacts. (A-H) Phase (A, C, E, G) and fluorescent membrane (B, D, F, H) images of MDCK cell island before stretch (A, B) and after full stretch (E, F) and those of MDCK vinculin KO cell island before stretch (C, D) and after full stretch (G, H). Full stretch here refers to 38% linear strain. Length scale bar is 10 µm. (I) Fraction of cell-cell contacts ruptured as a function of linear strain in percent. Each data point is the mean ± standard deviation for 306 MDCK cell-cell contacts and 378 MDCK vinculin KO cell-cell contacts, pooled from 2-4 large cell islands.

## Results and Discussion

Vinculin is known to be recruited to E-cadherin mediated adhesions at cell-cell contacts under the action of endogenous forces [19]. However, it is known that vinculin can also be recruited at cell-cell contacts in the absence of myosin mediated contractility [45]. Thus, we wanted to test whether vinculin is an essential element for sustaining high levels of endogenous force at cell-cell contacts or if its effect on endogenous inter-cellular tension is only marginal. To this end, we first generated a CRISPR knockout (KO) of vinculin in MDCK cells (fig. S1, fig. 1A,B). Immunofluorescence staining for vinculin marks focal adhesions in wildtype (WT) MDCK cells (fig. 1A), but this is not the case for MDCK vinculin KO cells (fig. 1B). Vinculin is an important focal adhesion protein as well [46-48], and previous reports [49, 50] have shown that the absence of vinculin decreases the traction force exerted by fibroblasts onto the ECM. Vinculin was also recently shown to be essential for high force transmission through focal adhesions in HeLa cells [51]. However, another study [52] reported that vinculin knockdown did not significantly decrease traction forces exerted by mesenchymal stem cells. Therefore, we first characterized our epithelial cells lacking vinculin (MDCK vinculin KO) by measuring the traction forces exerted by these cells on collagen-I coated soft substrates. The soft substrates we used were soft silicone (Qgel 300) of shear storage modulus 2.9 ± 1.2 kPa as determined using shear rheology (Fig. S2). Using traction force microscopy (see Methods), we found that vinculin KO cells exerted significantly lesser traction than WT cells (fig. 1C-G). The strain energy (which is a convenient proxy for the overall levels of traction forces exerted [53]) was 10.5 ± 6.1 fJ for vinculin KO versus 22.2 ± 16.3 fJ for WT (p < 0.05).

We then asked if the absence of vinculin affects the level of endogenous force transmitted via cell-cell contacts. To answer this, we used the Traction Force Imbalance Method (TFIM) [39]. Unlike an isolated cell, where the vector sum of traction forces vanishes (within experimental error), for each cell within a cell pair, the vector sum of traction forces is not balanced as such, but this imbalance in traction force corresponds to the inter-cellular force that is required for physical force balance for each cell in the cell pair. TFIM has been previously used to measure inter-cellular forces within endothelial cell pairs [54] and epithelial cell pairs undergoing dynamic cell rearrangements [55] and epithelial cell sheets [56]. We measured the traction forces for WT (fig. 2A,B) and vinculin KO cell pairs (fig. 2C,D) and then used TFIM to determine the inter-cellular force in WT and vinculin KO cell pairs (fig. 2E). We found that the cell-cell tension was significantly less for vinculin KO cell pairs than that for WT cell pairs (fig. 2E) – 23 ± 12 nN for vinculin KO cell-cell contacts versus 51 ± 24 nN for WT cell-cell contacts (p < 0.001). Thus, the absence of vinculin precludes cells from exerting high levels of endogenous tension through cell-cell contacts. Despite the absence of vinculin, cell-cell contacts in MDCK vinculin KO cell monolayers show the presence of actin, E-cadherin (fig. S3) and α-catenin (fig. S4).

Since it is α-catenin that is considered to be the primary recruiter of vinculin to E-cadherin adhesions, we surmised that α-catenin would be at least as important as vinculin in force transmission through cell-cell contacts. The centrality of α-catenin in potential force transmission pathways of E-cadherin to F-actin via many candidate proteins such as afadin, EPLIN and ZO-1, among others, also suggested that it would be a critical component of the effective mechanical pathway at cell-cell contacts. We thus expected the absence of α-catenin to decrease the endogenous forces at cell-cell contacts even more severely than the absence of vinculin. To test this, we generated an MDCK α-catenin KO cell line (fig. S1). We noticed that cell-cell contacts in α-catenin KO cell monolayers show the localization of both actin and E-cadherin (fig. S3). We then measured traction forces for cell pairs (fig. 3A,B) and then used TFIM to determine the endogenous force transmitted in cell-cell contacts within α-catenin KO cell pairs. To our surprise, we found that the inter-cellular tension for α-catenin KO cell-cell contacts was 39 ± 23 nN – not (statistically) significantly lesser than that for WT contacts (fig. 3E). However, our results are consistent with more qualitative laser ablation results – α-catenin knockdown (KD) was previously shown [21] to cause only a minor decrease in cell-cell tension as assessed by the retraction of the ablated ends of cell-cell contacts. α-catenin KD cells were also shown [37] to exert only slightly reduced traction forces on E-cadherin-coated substrates compared to WT cells.

Since vinculin is just one of many potential α-catenin binding partners that can transmit force to E-cadherin adhesion via the actin cytoskeleton, we wanted to test the specific role of the α-catenin-vinculin interaction. We exogenously expressed α-catenin lacking the vinculin binding site (VBS) (α-catenin DVBS) in MDCK α-catenin KO cells. We then measured the traction forces for MDCK α-catenin DVBS cell pairs (fig. 3C,D) and then used TFIM to determine the inter-cellular force. The inter-cellular force for MDCK α-catenin DVBS cell-cell contacts was 36 ± 20 nN – not (statistically) significantly lesser than that for WT contacts, similar to α-catenin KO cell-cell contacts (fig. 3E). We further tested whether the absence of both α-catenin and vinculin would drive down the inter-cellular tension lower than what we observed with the vinculin KO cells. Thus, we generated double KO cells where both α-catenin and vinculin were knocked out (fig. S1). Cell-cell contacts in monolayers of these double KO cells showed the absence of vinculin and α-catenin as expected (fig. S4), but still showed the localization of actin and E-cadherin (fig. S3). We then measured the traction forces for MDCK α-catenin-vinculin double KO cell pairs (fig. 4A,B) and then used TFIM to determine the inter-cellular force (fig. 4C). The cell-cell tension at MDCK α-catenin-vinculin double KO contacts was 21 ± 15 nN – similar to that for vinculin KO contacts and significantly less than that for WT contacts.

This result is consistent with the previously reported observation [37] that cells expressing α-catenin DVBS exert similar traction forces on E-cadherin-coated substrates as cells lacking α-catenin. Thus, it is vinculin rather than α-catenin that is essential for transmitting high endogenous forces at cell-cell contacts. While we used α-catenin KO as an experimental tool, it has been suggested that the decoupling of α-catenin from the E-cadherin-β-catenin complex may be physiologically relevant in cadherin junction disassembly in some contexts [16].

Given the essential role that we found for vinculin in the exertion of high endogenous inter-cellular forces, we wanted to know if vinculin also performs a similar essential role in protecting cell-cell contact integrity under mechanical challenges. In order to test this role for vinculin at cell-cell contacts, we wanted to use a method that can directly test the integrity of lateral cell-cell contacts between epithelial cells under external stretch. We plated either MDCK cells (fig. 5A,B) or MDCK vinculin KO cells (fig. 5C,D) on collagen-coated silicone sheets, and subjected the epithelial cell islands to large external stretch over a duration of a few minutes. We found that both MDCK cell islands (fig. 5E,F) and MDCK vinculin KO cell islands (fig. 5G,H) predominantly remained adhered to the substrate. Fortuitously, this allowed us to now assess the effect of the absence of vinculin on the integrity of cell-cell contacts. We found that cell-cell contacts ruptured over time within both MDCK (fig. 5E,F) and MDCK vinculin KO (fig. 5G,H) cell islands. We used a fluorescent live cell plasma membrane stain (CellBrite) to monitor what fraction of cell-cell contacts ruptured as a function of time. We found that MDCK vinculin KO cell-cell contacts ruptured at over twice the rate as that of MDCK cell-cell contacts (fig. 5I). Thus, vinculin is not only essential for high endogenous force transmission, but also for maintaining cell-cell contact integrity under high external forces. These results are consistent with previous reports of a mechanoprotective role for vinculin at E-cadherin adhesions, suggested by its recruitment to sites of forces exerted via E-cad-beads [32, 57]. Our results are also consistent with prior results using suspended doublets [58], detached cell sheets [36] and E-cadherin-coated substrates [59] that indicated a role for vinculin in maintaining cell-cell contact integrity. Our results with cell-cell contacts also complements the known role for vinculin in maintaining the integrity of cell-ECM contacts under cell-generated tension [60].

## Conclusion

Force transmission through epithelial cell-cell contacts plays a pivotal role in dynamic events during morphogenesis and adult tissue repair. In this report, we show that vinculin is essential for transmitting high levels of endogenous force through cell-cell contacts. Our results not only suggest that the α-catenin-vinculin complex is not necessary for transmitting high endogenous tension through cell-cell contacts, but also that α-catenin’s interaction with other proteins like afadin, EPLIN or ZO-1 is not essential for transmitting high inter-cellular tension. The α-catenin-vinculin may thus be just one of many active mechanical links for transmitting forces through cell-cell contacts. Through what other links may high levels of cell-generated forces be transmitted from the actomyosin apparatus to E-cadherin adhesions at cell-cell contacts? Our results are consistent with previously proposed interactions such as β-catenin-vinculin playing a mechanical role [21]. In fact, vinculin and α-catenin bind to the same N-terminal region of β-catenin [59, 61]. There is also evidence for the β-catenin-vinculin interaction in cancer cells that lack α-catenin [61] and for an α-catenin-independent means by which β-catenin can couple to the actin cytoskeleton [62]. However, the nanoscale positioning of vinculin is a bit displaced from β-catenin in α-catenin KD cells [21], suggesting that other intermediate molecular linkers may play a role. Vinculin can also be recruited to epithelial cell-cell contacts in a myosin VI dependent manner [20]. In any case, our results suggest that vinculin plays a key role at cell-cell contacts in addition to its established role of being recruited to α-catenin under specific force inputs. Vinculin’s ability to support high junctional tension as well as high contact strength, as shown here, is consistent with its essential role not only at cell-cell contacts in epithelia, but also other tissues undergoing dynamic events [63], including those like cardiac tissues [64], where endogenous forces reach even higher values. It is likely that high force transmission through vinculin enables the enhanced adhesion strength of cell-cell contacts, reminiscent of force coupled stabilization reported at focal adhesions [65, 66]. Vinculin’s key mechanical role at cell-cell contacts may also potentially explain why a bacterial pathogen has evolved to specifically bind to it to reduce cell-cell tension and promote its spread from cell to cell [67].

## Acknowledgements

We thank U. Schwarz for B. Sabass for the traction stress reconstruction script. The web app PlotsOfData [68] was used to render some of the plots.

## Author contributions

V.M. designed research; M.M., S.D., I.F, C.L, C.M, and J.I.C. performed research; D.C. contributed new reagents or analytic tools, M.M. and V.M. analyzed data; M.M. and V.M. wrote the paper.

