## Supplementary Figures for "Vinculin is Essential For Sustaining Normal Levels of Endogenous Force Transmission at Cell-Cell Contacts"

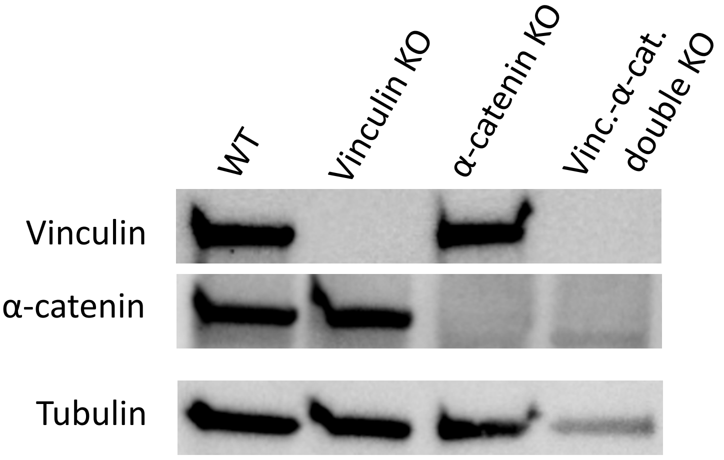


**Figure S1**. Western blot image of lysed MDCK (WT), MDCK vinculin KO, MDCK α-catenin KO and MDCK vinculin-α-catenin double KO cells, probed for vinculin, α-catenin and tubulin (loading control). Note that the loading level of the last lane (MDCK vinculin-α-catenin double KO lane) is notably lesser, however there clearly is no band corresponding to vinculin or α-catenin.

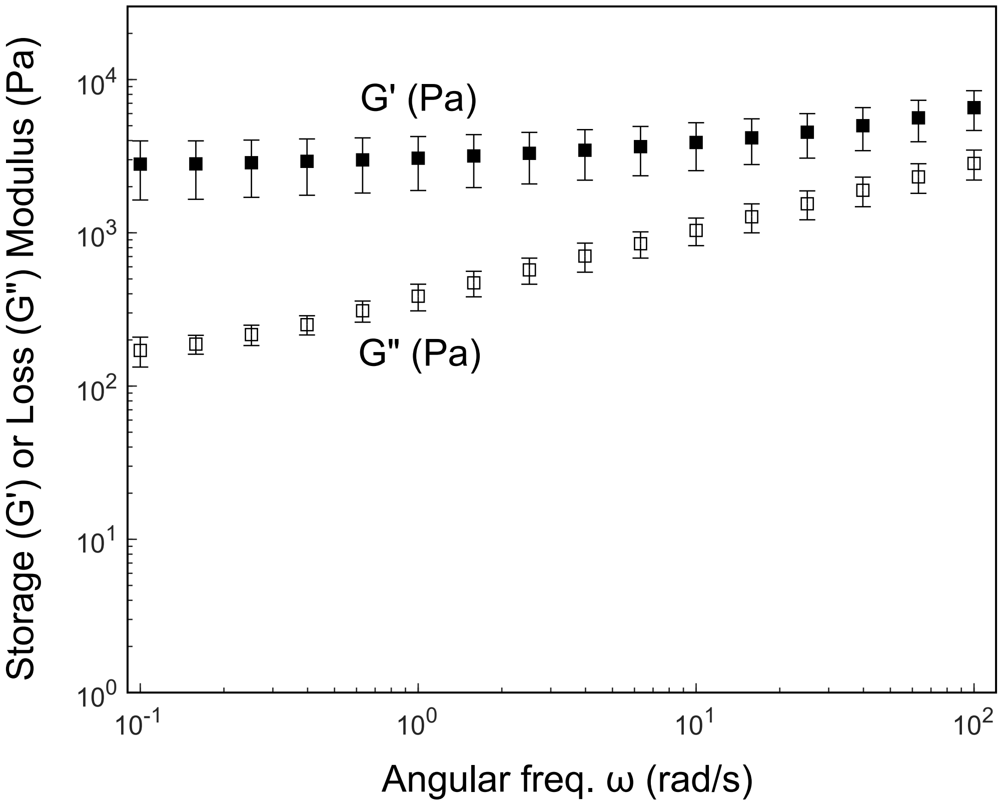


**Figure S2**. Shear rheological characterization of soft silicone Qgel (A:B = 1:2.2). Shear storage and loss moduli are plotted as mean ± standard deviation for each angular frequency of oscillation.

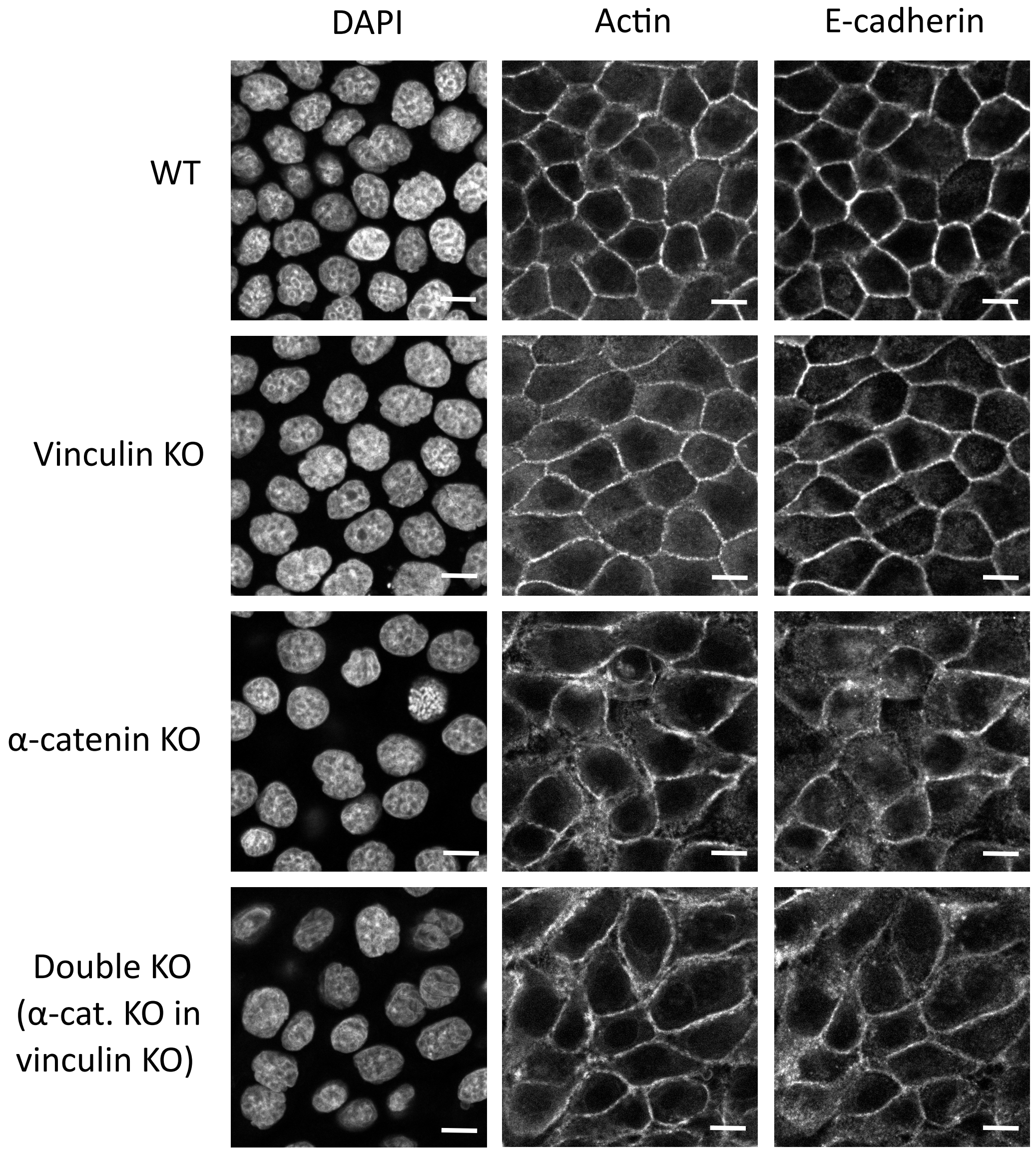


**Figure S3**. Immunofluorescence images of MDCK (WT), MDCK vinculin KO, MDCK α-catenin KO and MDCK vinculin-α-catenin double KO cells, stained for the nucleus (DAPI), actin and E-cadherin. Scale bar is 10 μm.

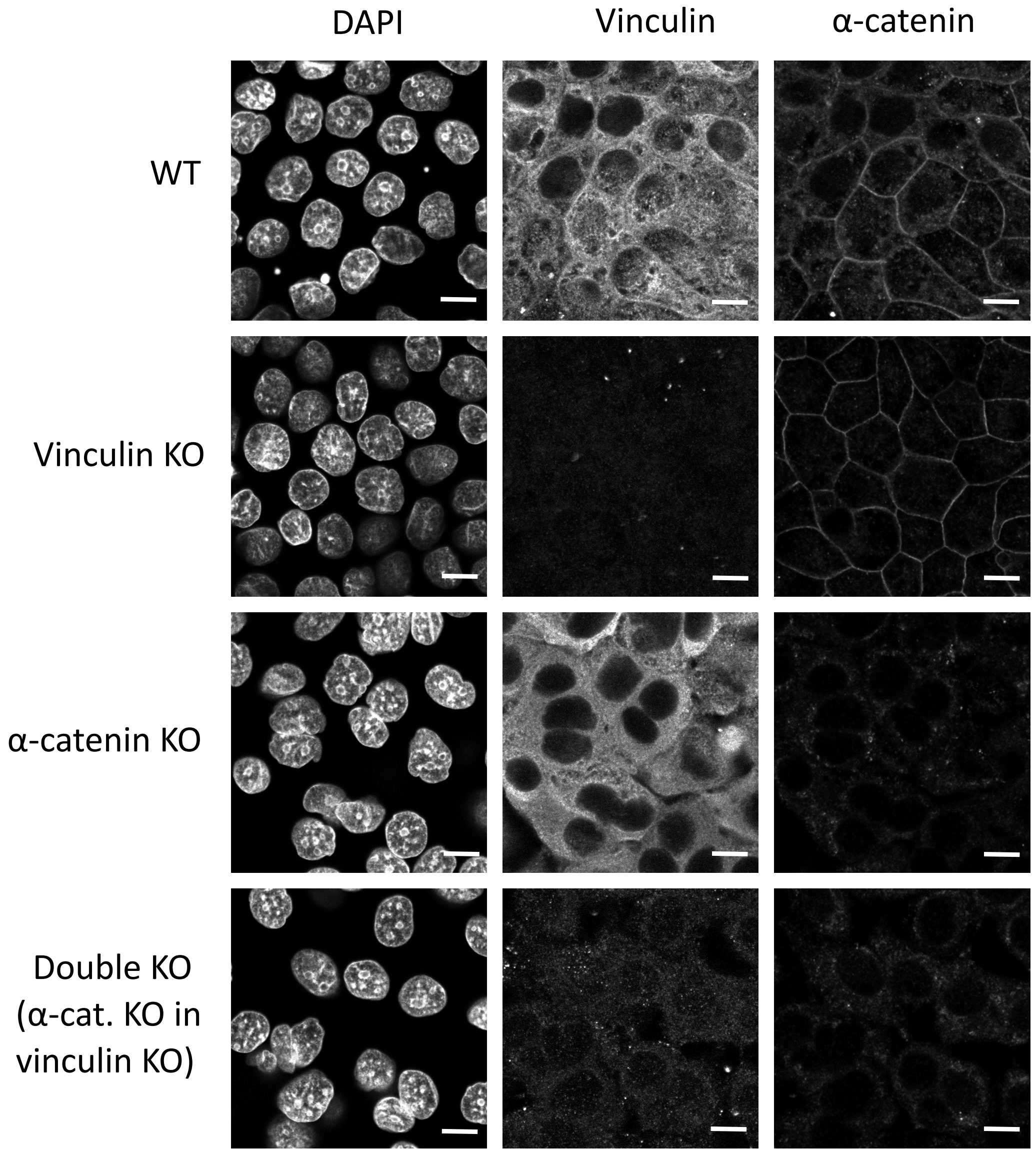


**Figure S4**. Immunofluorescence images of MDCK (WT), MDCK vinculin KO, MDCK α-catenin KO and MDCK vinculin-α-catenin double KO cells, stained for the nucleus (DAPI), vinculin and α-catenin. Scale bar is 10 μm.
